# SpheroMap Cytometry: a novel spatial flow cytometry approach to evaluate immune response in PDAC spheroids

**DOI:** 10.1101/2025.08.08.669354

**Authors:** Sara E. Castro-Silva, Camila C. O. M. Bonaldo, Patricia V. B. Palma, Pedro Luiz Porfirio Xavier, Raphael Lucas More, Maristela D Orellana, Sâmia R. Caruso, Rodrigo A. Panepucci

## Abstract

Pancreatic ductal adenocarcinoma (PDAC) is characterized by a dense stroma that contributes, along with hypoxia, to immune exclusion and therapeutic resistance. Here, we introduce SpheroMap Cytometry, an innovative spatial flow cytometry platform designed to preserve and quantify the relative localization of immune cells from 3D spheroid models. CAPAN-1 pancreatic tumor cells and either HS-5 bone marrow stromal cells or umbilical cord-derived mesenchymal stromal cells (UC-MSCs) co-cultured in ultra-low-adhesion 96-well plates. PBMC cells (activated or non-activated) were added to infiltrate pre-formed tumor–stromal/mesenchymal spheroids. Then after 48 hours post-seeding and 24 hours post-PBMC incubation, spheroids were incubated with Image-iT Green Hypoxia to mark hypoxic cores, and after 72 hours spheroids were dissociated and stained with CD3, CD4, CD8, CD25 and CD127 antibodies. SpheroMap Cytometry revealed that non-activated immune cells infiltrated spheroids with a distinct pattern between normoxic and hypoxic regions, with an enrichment of CD4^+^ over CD8^+^ cells in the hypoxic core and a higher proportion of CD4^+^CD25^+^CD127^−^ Treg-like cells. Pre-activation of PBMCs enhanced CD8^+^ cell infiltration into the hypoxic region and increased CD25^+^/CD25^high^/CD127^−^ Tregs in both compartments, indicating that T-cell activation, while facilitating CD8^+^ cell entry into hypoxic zones, may also promote immunosuppressive populations that impair cytotoxic function. Notably, the effects we observed from the infiltration of pre-activated lymphocytes into tumor–stromal spheroids were not seen in spheroids formed with tumor–mesenchymal cells, which may reflect the immunosuppressive role of umbilical-cord mesenchymal cells. Our findings demonstrate that hypoxia modulates T-cell infiltration and activation, underscoring the critical interplay between oxygen gradients and immune evasion in PDAC. SpheroMap Cytometry provides a scalable, high-resolution method to dissect the spatial immunophenotype of lymphocytes in PDAC models, overcoming limitations of static imaging and conventional flow cytometry, making it an ideal platform for testing therapies aimed at overcoming immune exclusion.

## 1 Introduction

Pancreatic ductal adenocarcinoma (PDAC) carries one of the worst prognoses among solid malignancies due to nonspecific symptoms, late diagnoses, and therapy resistance, contributing to a five-year survival rate below 11% ^1^. PDAC has a characteristically immune-excluded profile, characterized by accumulation of immune cells confined to the periphery of cancer cell nests, wich suggests effective long-distance recruitment, but infiltration is blocked by physical and/or functional barriers.

Through activation of the Hypoxia-Inducible Factor (HIF), hypoxia can affect the recruitment, infiltration, survival, activation, and function of various immune cells; for example, by promoting an immunosuppressive M2 phenotype in myeloid and stromal cells and driving the generation of Tregs, thereby directly impacting the immune landscape and antitumor response. Hypoxia–HIF-1α signaling suppresses both innate and adaptive immunity through multiple mechanisms, inducing the expression of immunosuppressive factors (TGF-β, IL-10, VEGF, PGE_2_) and immune checkpoint molecules (e.g., PD-1 and PD-L1/2), and inhibiting tumor antigen presentation via MHC-I. Nutrient deprivation and the accumulation of toxic catabolic byproducts (such as lactate) severely impair effector T-cell functions, inhibiting the cytotoxic potential of CD8^+^ T lymphocytes and NK cells while, conversely, conferring a metabolic advantage to Tregs. HIF-1α also induces a shift from IFN-γ–producing Th1 cells to IL-10–producing Th2 cells and upregulates inhibitory checkpoint receptors on activated T cells. Moreover, NK cells cultured under hypoxia are functionally impaired, and intratumoral NK cells exhibit hypoxia signatures and are enriched for a CD73^+^ “regulatory NK” subpopulation (expressing checkpoints such as LAG3, VISTA, PD-1, and PD-L1), which produce IL-10 (via STAT3) and suppress CD4^+^ T-cell proliferation as well as IFN-γ production.

In recent years, it has become clear that the tumor “immune context” prior to therapy (i.e., the quantitative and qualitative landscape of infiltrating immune cells) directly impacts the prognosis of cancer patients. In this context, understanding how hypoxia affects the distribution, function, and survival of immune cells within the tumor microenvironment represents an important step in the development of new therapies. Considering these challenges, this study proposes to evaluate how hypoxia influences the infiltration and distribution of immune cells in spheroid models, through a novel Spatial Flow Cytometry approach (SpheroMap Cytometry) thereby providing a significant methodological advance for the field of translational immuno-oncology.

## 2 Methodology

### 2.1 Cell Lines and Culture

CAPAN-1 pancreatic cancer cells (BCRJ, 265) were kindly provided by Dr. Pedro Luiz Porfirio Xavier, collaborating researcher in the Laboratory of Comparative and Translational Oncology (LOCT) at FZEA/USP. HS-5 immortalized stromal cells (ATCC CRL-3611) and umbilical-cord-derived mesenchymal cells (UC-MSCS) were generously supplied by Dr. Sara Teresinha Olalla Saad, Full Professor at the State University of Campinas, and Dr. Maristela Delgado Orellana, researcher at the Hemocenter of Ribeirão Preto, respectively.

The CAPAN-1, HS-5, and primary UC-MSCS cell lines were cultured in IMDM, DMEM, and α-MEM, respectively, each supplemented with 10% fetal bovine serum, 10% penicillin/streptomycin, and HEPES. Cultures were maintained in a 37 °C incubator with 5% CO_2_ and 85% humidity.

### 2.2 Processing and Activation of Peripheral Blood Mononuclear Cells (PBMCs)

For analysis of lymphocyte infiltration into the developed spheroid model, peripheral blood mononuclear cells (PBMCs) from healthy donors were obtained from the Hemocenter of Ribeirão Preto following written informed consent. PBMCs were isolated by density gradient centrifugation using Ficoll-Paque. After isolation, PBMCs were labeled with Cell Proliferation Dye eFluor 450 (Invitrogen™ 65-0842-85) at a final concentration of 10 µM, resuspended in RPMI medium, and incubated. On the day following labeling, a portion of the cells was activated for 24 hours with lymphocyte stimulators ionomycin (Sigma I0634; 1 µg/mL) and PMA (phorbol 12-myristate 13-acetate, Sigma P1585; 50 ng/mL). After this 24-hour period, the PBMCs (activated or non-activated) were washed and resuspended in RPMI

### 2.3 Spheroid Establishment

A total of 20,000 CAPAN-1 and HS-5 or CAPAN-1 and UC-MSCS cells at a 1:4 tumor-to-stromal/mesenchymal ratio were co-plated in ultra-low-adhesion (ULA) 96-well plates (Thermo Scientific 174115). After 24 hours of incubation, 100,000 PBMC cells pre-labeled with Cell Proliferation Dye eFluor 450 (activated or non-activated) were added to infiltrate pre-formed, unlabeled tumor–stromal/mesenchymal spheroids for 48 hours.

### 2.4 Spatial Flow Cytometry (SpheroMap Cytometry)

Given the limitations of conventional flow cytometry and microscopy approaches, we developed a novel spatial flow cytometry method (SpheroMap Cytometry) to enable more detailed immunophenotypic and spatial analysis of the different cell populations within spheroids. Our approach is based on the unique properties of the Image-iT Green Hypoxia reagent, which accumulates intracellular fluorescence specifically in cells located within the hypoxic core of the spheroid. Importantly, this fluorescent signal is retained even after enzymatic dissociation of the spheroids, thus allowing for quantitative immunophenotypic analysis of the spatial distribution of distinct cellular subpopulations within the hypoxic (central) and normoxic (peripheral) compartments of the spheroid.

#### 2.4.1 Distribution of Lymphocyte Populations in Spheroid Compartments

After 48 hours post-seeding and 24 hours post-PBMC incubation, half of the medium was replaced with medium containing Image-iT Green Hypoxia (at a final concentration of 1.25 µM). Subsequently (at 72 hours post-seeding and 48 hours post-PBMC incubation), spheroids were collected, transferred to 15 mL centrifuge tubes (8 spheroids per tube), washed, and incubated in 2 mL of 0.05% Trypsin-EDTA (Gibco 15400054) for 10 to 15 minutes to allow enzymatic and mechanical dissociation (using a 200 µL pipette tip). Additionally, PBMCs that remained at the bottom of the well and did not infiltrate the spheroids (non-infiltrating PBMCs) were collected and separated for analysis (Fig. 1)

**Fig. 1.**
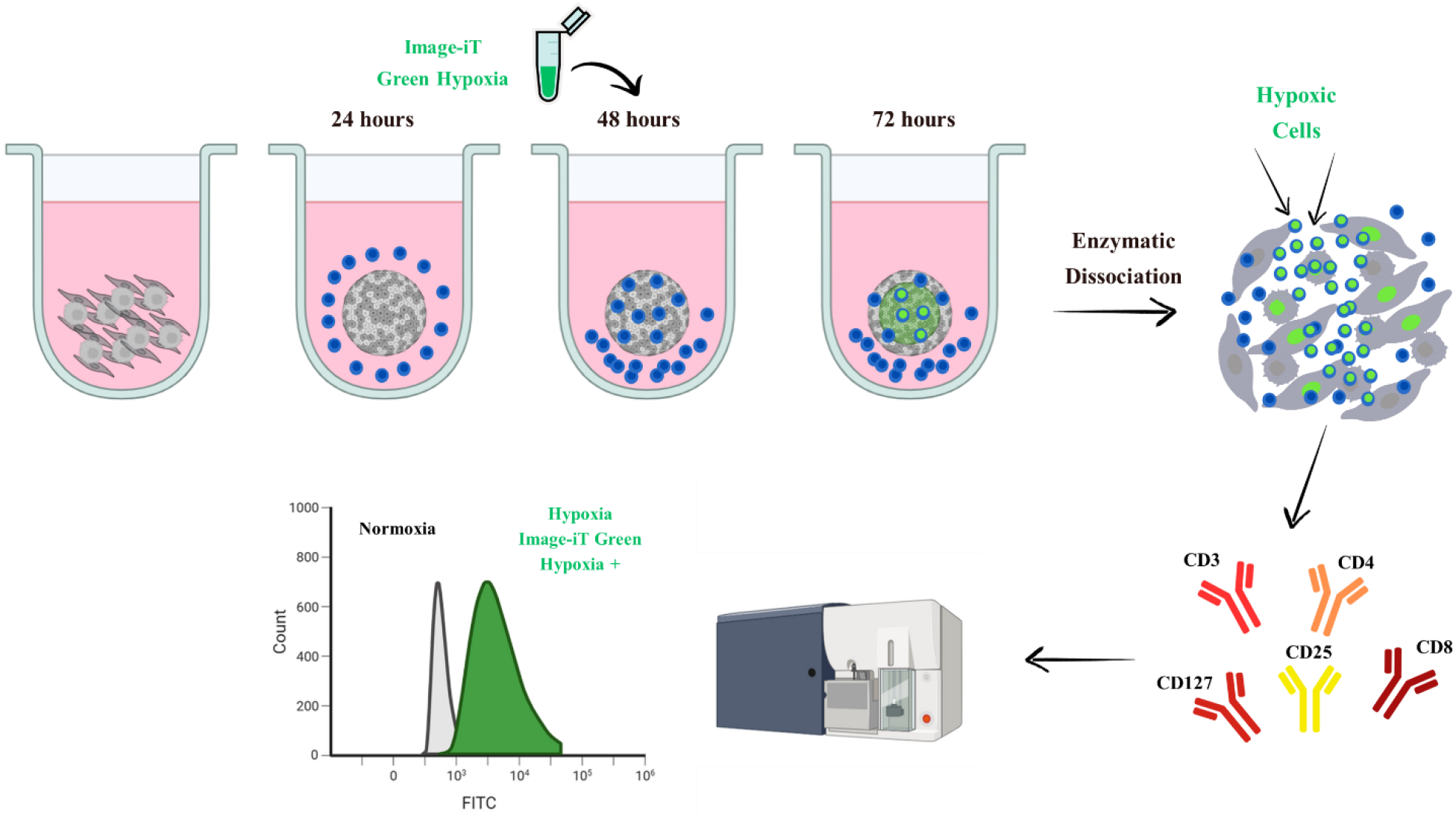
Schematic of the SpheroMap Cytometry methodology. A total of 20,000 tumor and stromal/mesenchymal cells were co-plated in U-bottom ultra-low attachment plates at a 1:4 tumor-to-stromal ratio. After 24 hours of incubation, 100,000 PBMC cells pre-labeled with Cell Proliferation Dye eFluor 450 (activated or non-activated) were added to infiltrate pre-formed tumor–stromal/mesenchymal spheroids. Then after 48 hours post-seeding and 24 hours post-PBMC incubation, the Image-iT Green Hypoxia (1.25 µM) reagent was added to wells containing the spheroids. Intracellular fluorescence accumulates in cells occupying the hypoxic core of the spheroid and, importantly, is retained even after enzymatic dissociation, enabling immunophenotypic analysis of cells that inhabited the hypoxic region. 72 hours after initial seeding, spheroids were enzymatically dissociated, stained with CD3, CD4, CD8, CD25 and CD127 antibodies, and subjected to analysis.

The dissociated cells were then washed with RPMI medium, centrifuged, and resuspended in PBS + 2% FBS, followed by incubation for 30 minutes at 4 °C with the following antibodies: APC Mouse Anti-Human CD3 (Clone UCHT1, BD Pharmingen™ 555335), BV605 Mouse Anti-Human CD4 (Clone RPA-T4, BD OptiBuild™ 740563), PE-Cy™7 Mouse Anti-Human CD8 (Clone RPA-T8, BD Pharmingen™ 560917), PE Mouse Anti-Human CD25 (Clone M-A251, BD Pharmingen™ 555432), and BB700 Mouse Anti-Human CD127 (Clone HIL-7R-M21, BD Horizon™ 566398). After the incubation period, cells were washed, centrifuged, and resuspended in PBS + 2% FBS in flow cytometry tubes and then subjected to analysis.

### 2.5 Analysis of SpheroMap Cytometry Data

SpheroMap Cytometry data were acquired using a BD FACSLyric™ Clinical Flow Cytometry System, then exported and analyzed using FlowJo software v.10.9.0. Data obtained in FlowJo were subsequently exported to GraphPad Prism 9 for graph generation and statistical analysis. The statistical test used was ordinary two-way ANOVA followed by Sidak’s multiple comparisons test.

## 3 Results

### 3.1 Distribution of immune populations in tumor-stromal spheroids

To understand how PBMC activation and the spheroid TME influence lymphocyte phenotype and infiltration, we performed two comparative analyses using SpheroMap Cytometry: (i) analysis of PBMCs that infiltrated the spheroids (both activated and non-activated), characterizing their localization within hypoxic or normoxic compartments, and (ii) immunophenotypic characterization of the recovered PBMCs that did not infiltrate the spheroids (activated and non-activated). As described in the literature and observed in our results, activation with PMA and ionomycin leads to a decrease in CD4 expression. Therefore, we used the marker for CD8 T lymphocytes as a control, classifying CD3^+^CD8^−^ cells as CD4 T lymphocytes ^2-4^.

PBMCs analyzed prior to spheroid incubation displayed a typical lymphocyte profile, with an average CD4^+^:CD8^+^ ratio of 2:1 within the CD3^+^ T-cell population, and a slight enrichment of CD4^+^ cells under the activated condition. In PBMCs recovered from spheroids formed with CAPAN-1 and HS-5 cells, when comparing the activated condition to the non-activated one, we observed that the normoxic periphery exhibited a reduction in CD3^+^ lymphocytes (20.73% vs. 33.80%), likely due to increased infiltration of this population into the hypoxic compartment under activation. The proportion of CD3^+^CD4^+^ cells also increased (68.37% vs. 55.57%), accompanied by elevated CD4^+^CD25^+^ expression (48.40% vs. 4.98%) and CD4^+^CD25^high^ cells (8.92% vs. 0.40%). Regulatory T-cell frequency (CD4^+^CD25^+^CD127^−^) also rose in normoxia when comparing activated to non-activated conditions (28.07% vs. 1.85%) (Fig. 2A-E). The cytotoxic CD8^+^ population decreased significantly (29.77% vs. 43.40%), again reflecting enhanced CD8^+^ infiltration into hypoxic regions under activation. However, we saw significant increases in CD8^+^CD25^+^ (68.67% vs. 8.97%) and CD8^+^CD25^high^ (14.50% vs. 0.12%) expression. In the hypoxic core, pre-activation of PBMCs promoted greater CD3^+^ lymphocyte infiltration (79.27% vs. 66.20%). The proportion of CD3^+^CD4^+^ cells decreased (51.73% vs. 74.13%), but CD4^+^CD25^+^ expression rose markedly (92.13% vs. 74.30%), as did CD4^+^CD25^high^ cells (44.87% vs. 9.51%) and Treg frequency (CD4^+^CD25^+^CD127^−^), which more than doubled (48.93% vs. 18.30%). The cytotoxic CD8^+^ population also increased significantly (46.13% vs. 24.80%), with substantial gains in CD8^+^CD25^+^ (98.73% vs. 62.20%) and CD8^+^CD25^high^ (39.77% vs. 9.71%) expression (Fig. 2A-E).

**Fig. 2.**
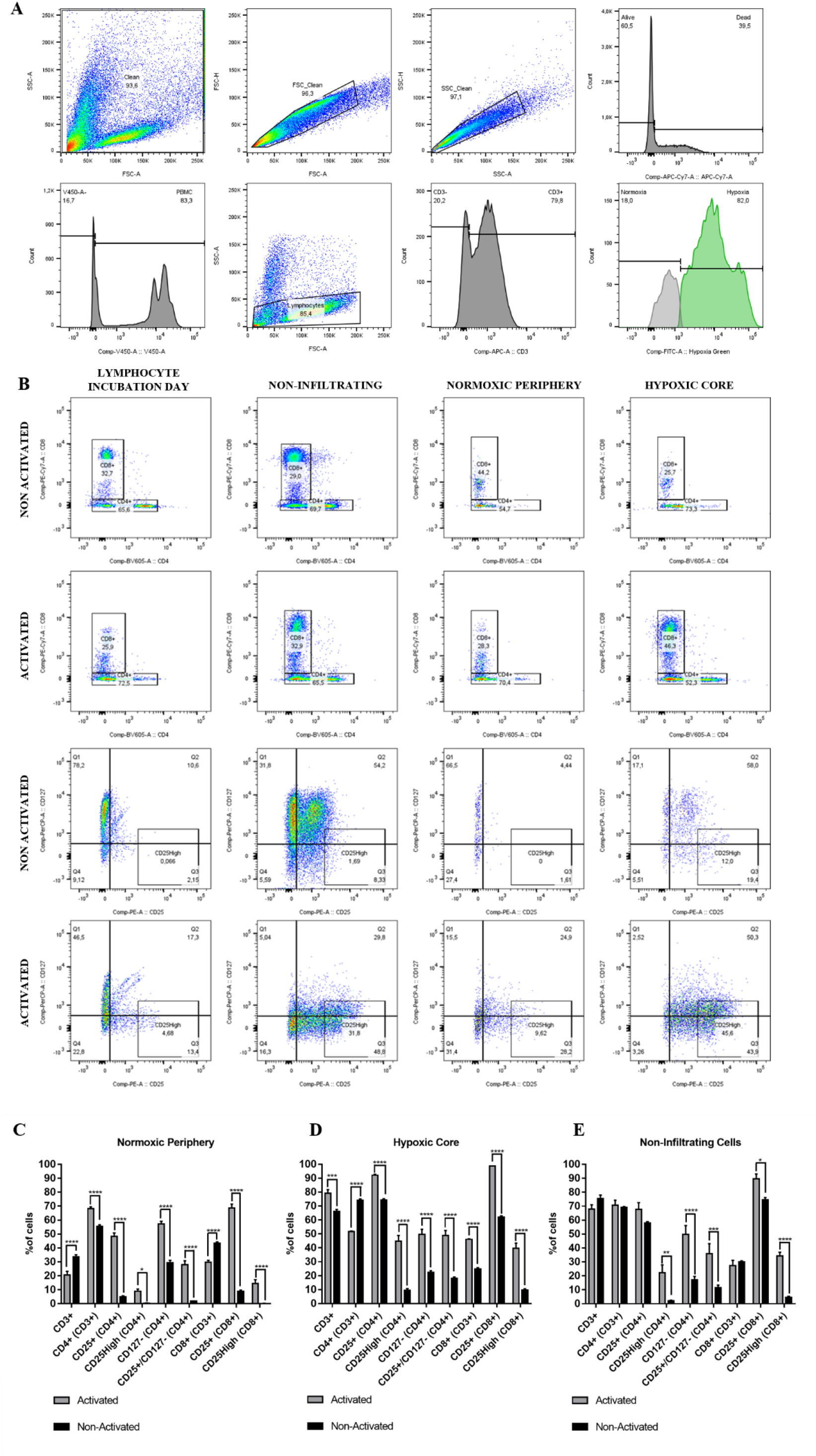
T-cell subsets distribution in CAPAN-1+HS-5 tumor–stromal spheroids and non-infiltrating PBMCs. **(A-B)** 20.000 tumor and stromal cells were co-seeded in U-bottom ultra-low-attachment plates at a 1:4 ratio. After 24 h, 100,000 pre-labeled PBMCs (activated or not) were added to infiltrate the formed spheroids. At 48 h post-seeding (24 h post-PBMC addition), Image-iT Green Hypoxia (1.25 µM) was applied, and at 72 h, spheroids were enzymatically dissociated and stained for CD3, CD4, CD8, CD25, and CD127. Flow cytometry gating sequentially excluded debris, singlets, and dead cells, identified CP450^+^ PBMCs versus CP450^−^ tumor/stromal cells, and distinguished lymphocyte subsets under hypoxia (Hypoxia Green^+^) and normoxia (Hypoxia Green^−^). In each compartment, percentages of CD3^+^, CD4^+^, CD8^+^, CD25^+^, CD127^−^, and CD25^+^CD127^−^ T cells were quantified. **(C-E)** Quantification of non-infiltrating PBMCs remaining in the culture supernatant. Statistical analysis was performed using ordinary two-way ANOVA followed by Sidak’s multiple comparisons test. *p < 0.05, **p < 0.01, ***p < 0.001, ****p < 0.0001.

Non-infiltrating PBMCs maintained similar CD3^+^, CD3^+^CD4^+^, and CD3^+^CD8^+^ profiles under both activated and non-activated conditions. However, in the activated condition we observed significant increases in CD4^+^CD25^high^ (22.40% vs. 2.17%), CD4^+^CD25^+^CD127^−^ (35.87% vs. 11.61%), CD8^+^CD25^+^ (89.63% vs. 74.50%), and CD8^+^CD25^high^ (34.33% vs. 4.58%) compared to the non-activated condition (Fig. 2A-E).

### 3.2 Distribution of immune populations in tumor-mesenchymal spheroids

Lymphocyte infiltration into spheroids formed with CAPAN-1 and UC-MSCS cells exhibited some differences compared to results from CAPAN-1 + HS-5 spheroids. When comparing the activated condition to the non-activated one, we observed a reduction in CD3^+^ lymphocytes in the normoxic periphery (25.53% vs. 44.93%), likely due to increased infiltration into the hypoxic compartment under activation. The proportion of CD3^+^CD4^+^ cells also increased (82.07% vs. 61.23%), along with elevated CD4^+^CD127^−^ expression (62.63% vs. 33.07%). The cytotoxic CD8^+^ population decreased significantly (17.30% vs. 37.93%), though we saw a significant rise in CD8^+^CD25^+^ expression (31.20% vs. 8.19%). In the hypoxic core, pre-activation of PBMCs promoted greater CD3^+^ lymphocyte infiltration (74.47% vs. 55.07%) but did not alter the proportions of CD3^+^CD4^+^ and CD3^+^CD8^+^ cells. However, the frequency of Tregs (CD4^+^CD25^+^CD127^−^) increased (31.17% vs. 20.07%), and CD8^+^CD25^+^ expression also rose (87.00% vs. 70.47%) (Fig. 3A-E).

**Fig. 3.**
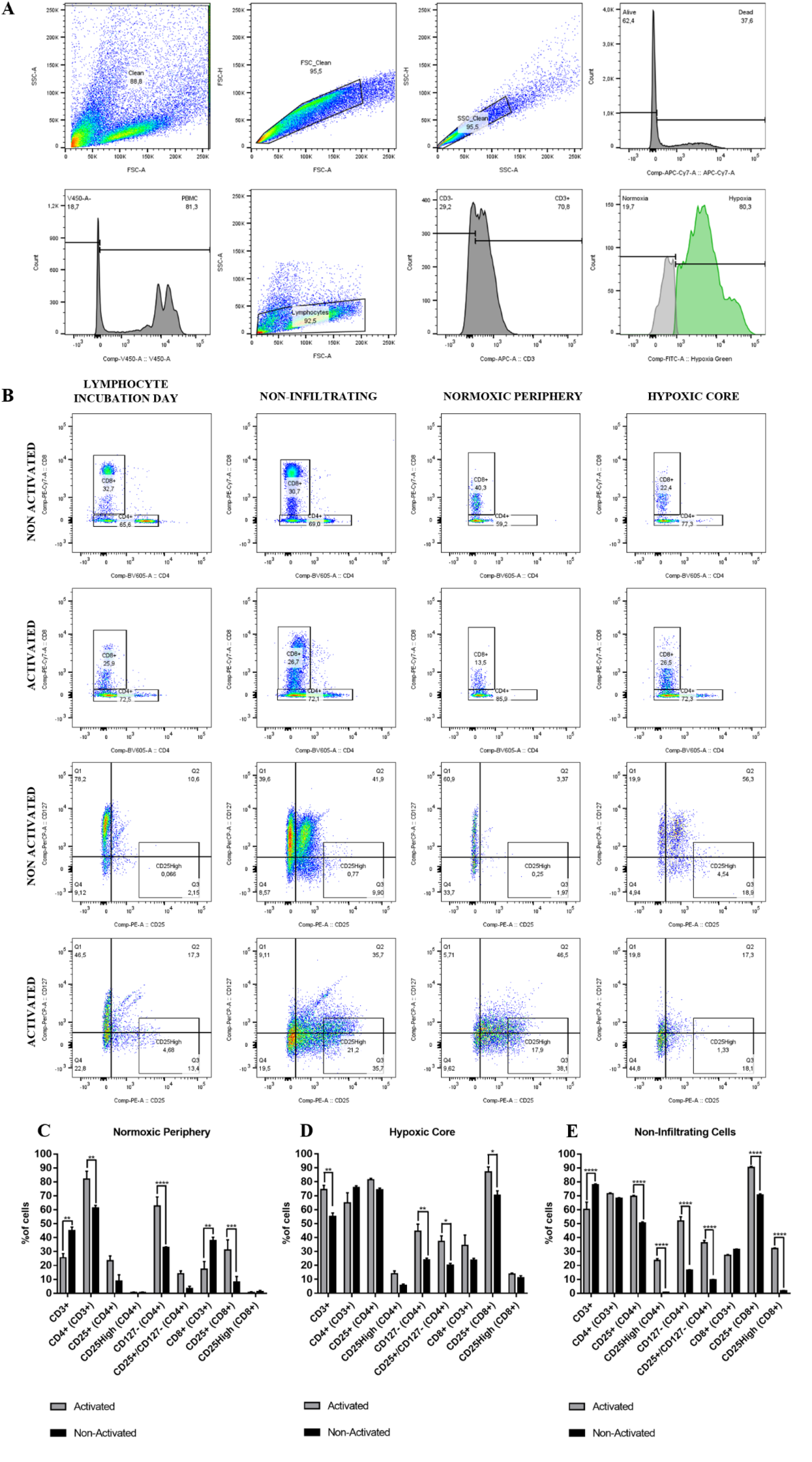
T-cell subsets distribution in CAPAN-1+UC-MSCS tumor–mesenchymal spheroids and non-infiltrating PBMCs. **(A-B)** 20.000 tumor and mesenchymal cells were co-seeded in U-bottom ultra-low-attachment plates at a 1:4 ratio. After 24 h, 100,000 pre-labeled PBMCs (activated or not) were added to infiltrate the formed spheroids. At 48 h post-seeding (24 h post-PBMC addition), Image-iT Green Hypoxia (1.25 µM) was applied, and at 72 h, spheroids were enzymatically dissociated and stained for CD3, CD4, CD8, CD25, and CD127. Flow cytometry gating sequentially excluded debris, singlets, and dead cells, identified CP450^+^ PBMCs versus CP450^−^ tumor/stromal cells, and distinguished lymphocyte subsets under hypoxia (Hypoxia Green^+^) and normoxia (Hypoxia Green^−^). In each compartment, percentages of CD3^+^, CD4^+^, CD8^+^, CD25^+^, CD127^−^, and CD25^+^CD127^−^ T cells were quantified. **(C-E)** Quantification of non-infiltrating PBMCs remaining in the culture supernatant. Statistical analysis was performed using ordinary two-way ANOVA followed by Sidak’s multiple comparisons test. *p < 0.05, **p < 0.01, ***p < 0.001, ****p < 0.0001.

Activated non-infiltrating PBMCs exhibited a reduction in the percentage of CD3^+^ lymphocytes (60.23% vs. 78.07%) compared to non-activated PBMCs, likely reflecting enhanced infiltration into the spheroids. No significant differences were observed between activated and non-activated conditions for CD3^+^CD4^+^ and CD3^+^CD8^+^ populations; however, activated PBMCs demonstrated significant increases in CD4^+^CD25^+^ expression (69.60% vs. 50.47%), CD4^+^CD25^high^ (23.53% vs. 0.84%), CD4^+^CD25^+^CD127^−^ (36.20% vs. 9.81%), CD8^+^CD25^+^ (90.43% vs. 70.60%), and CD8^+^CD25^high^ (32.23% vs. 1.86%) relative to non-activated PBMCs (Fig. 3A-E).

## 4 Discussion

PDAC is well known for its immunologically excluded profile, in which the dense stromal compartment plays a key role in establishing physical and functional barriers that limit immune cell access ^5-7^. Through activation of the Hypoxia-Inducible Factor (HIF), hypoxia directly influences immune cell infiltration and distribution. Hypoxia–HIF-1α signaling suppresses both innate and adaptive immunity through multiple mechanisms, inducing the expression of immunosuppressive factors (TGF-β, IL-10, VEGF, PGE_2_) and immune checkpoint molecules (e.g., PD-1 and PD-L1/2), while inhibiting tumor antigen presentation via MHC-I ^8, 9^. Nutrient deprivation and the accumulation of toxic catabolic byproducts, such as lactate, severely impair effector T-cell functions ^10, 11^, inhibiting the cytotoxic potential of CD8^+^ T lymphocytes ^12^ and, conversely, conferring a metabolic advantage to regulatory T cells (Tregs) ^13, 14^. In our model, infiltration of non-activated immune cells into the spheroid displayed a differential profile between the normoxic and hypoxic compartments, revealing enrichment of CD4^+^ cells over CD8^+^ cells in the hypoxic region, as well as increased proportions of CD4^+^CD25^+^ cells and CD127^−^ Treg-phenotype cells.

A study conducted by our research group, in which SpheroMap Cytometry was used to identify CAF distribution in the tumor microenvironment of PDAC spheroids and their CD73 expression, linked the hypoxic environment to a CD73^High^CD140B^+^ stromal immunophenotype associated with potential myCAFs. Hypoxia plays a well-established role in CAF reprogramming and activation by promoting tumor-cell secretion of TGF-β and PDGF, which favors differentiation of precursor cells into myCAFs. These myCAFs, in turn, contribute significantly to desmoplasia through extracellular matrix deposition, leading to immunosuppression by excluding immune cells ^15-20^. CAFs also dramatically shape the tumor immune landscape through secretion of immunomodulatory cytokines such as TGF-β, IL-6, CXCL1, and CXCL12. One direct consequence of tumor–CAF crosstalk via TGF-β signaling is a reduction in cytotoxic T cells accompanied by increased Treg infiltration. More specifically, myCAFs have been shown to exert a suppressive effect on CD8^+^ T cells ^19, 21-23^. As confirmed in the study conducted by our research group, CD73 is highly expressed in PDAC tumors, particularly in stromal cells ^24-27^; previous work has shown that CD73 overexpression is associated with reduced infiltration of CD8^+^ tumor-infiltrating lymphocytes and with Treg-mediated immunosuppression, since this mechanism depends on extracellular adenosine generated by CD73 ^28, 29^. In this context, our findings align with published data on the impact of the stromal component and hypoxia on immune-cell infiltration ^30-34^.

Interestingly, when PBMCs were activated prior to infiltration into the spheroids, we observed an overall increase in PBMC and lymphocyte infiltration in spheroids formed with tumor– stromal cells, most notably a greater infiltration of CD8^+^ cells into the hypoxic compartment. These effects were not seen in spheroids formed with tumor–mesenchymal cells, which may reflect the immunosuppressive role of umbilical-cord–derived mesenchymal cells. MSCs induce T-cell apoptosis and cell-cycle arrest through secretion of indoleamine 2,3-dioxygenase (IDO), prostaglandin E_2_ (PGE_2_), and transforming growth factor–β1 (TGF-β1), and they also suppress T-cell activation and proliferation by modulating T-cell phenotypes and increasing the frequency of regulatory T cells ^35-39^. Activation also increased the expression of CD25^+^/CD25^high^ cells and CD127^−^ Tregs in both spheroid compartments, demonstrating that T-cell pre-activation, while capable of overcoming hypoxia- and stroma-imposed barriers for CD8^+^ cells, can also promote immunosuppressive cell populations, thereby compromising cytotoxic T-cell function. This suggests that strategies to overcome the observed immunosuppression may lie in modulating hypoxia-triggered pathways—such as adenosinergic signaling—through targeting CD73 and CAFs, and more specifically, myCAFs.

## 5 Conclusion

The innovative technique developed by our group, SpheroMap Cytometry, proved to be a powerful and scalable tool for inferring the relative spatial localization of immune cells in PDAC spheroids, overcoming the limitations of conventional approaches such as static microscopy and traditional flow cytometry. Using this technique, we were able to demonstrate that hypoxia distinctly modulates the infiltration and activation of T-cell subsets—enriching for CD4^+^ cells in the absence of activation and for CD8^+^ cells and Tregs following pre-activation, especially within hypoxic cores.

These findings underscore the central role of hypoxic gradients in modulating cell behavior within the TME and highlight the importance of approaches that combine physiological relevance with high analytical capability. The platform developed in this study not only advances our understanding of immune-evasion mechanisms in PDAC but also represents a promising tool for preclinical screening of immunotherapies and combination treatments targeting the TME, with potential translational applications in experimental oncology.

## 6 Acknowledgements

This study was funded by the São Paulo Research Foundation (FAPESP) Brasil, process #2022/12856-6; the Brazilian Federal Agency for Support and Evaluation of Graduate Education (CAPES); and the National Council for Scientific and Technological Development (CNPQ).

